# Generation and initial characterization of in vivo knockout of tetherin/BST2 in chicken

**DOI:** 10.1101/2024.06.24.600038

**Authors:** Lenka Ungrová, Pavel Trefil, Jiří Plachý, Jitka Mucksová, Jiří Kalina, Markéta Reinišová, Sonja Härtle, Eliška Gáliková, Dana Kučerová, Veronika Krchlíková, Vladimír Pečenka, Vít Karafiát, Jiří Hejnar, Daniel Elleder

## Abstract

Tetherin/BST2 is an antiviral restriction factor initially described in mammals. It is active against multiple enveloped viruses at the budding phase, where it is able to physically link the budding virions to the virus-producing cell. We and others have previously identified tetherin orthologs in birds, and characterized the antiviral activity and interferon-inducibility of chicken tetherin. In this work, we have generated an *in vivo* model of tetherin absence in chicken by CRISPR/Cas9 modification of chicken primordial germ cells (PGC). The modified PGCs were transplanted into roosters with suppressed endogenous spermatogenesis, and transgenic (tetherin knockout) progeny was obtained by further crosses. The viability and phenotype of tetherin knockout animals did not differ from wild type chicken. In more detailed investigation, flow cytometry based differential white blood cell count revealed an increased number of heterophils in tetherin knockouts. Upon challenge with avian sarcoma and leukosis virus (ASLV), a prototypic avian retrovirus, we detected increase in viremia at days 6 and 13 post infection in tetherin knockout animals. The increased virus susceptibility is consistent with absence of antiviral tetherin. In summary, we introduce a new *in vivo* knockout model of chicken antiviral gene tetherin. These animals can be used in further characterizations of avian antiviral defenses and also to define thus far unknown physiological effects of tetherin in birds.

## 1. INTRODUCTION

Birds represent important hosts for a variety of pathogens, including viruses with large economic impact and zoonotic potential (Nabi et al. 2021). A major type of antiviral defense is executed by dedicated antiviral genes, also called antiviral restriction factors, whose products target various steps in viral replication cycle (Duggal and Emerman 2012). Tetherin (also known as bone marrow stromal antigen 2 [BST2] or cluster of differentiation 317 [CD317]) was originally identified as a surface protein expressed on terminally differentiated B-cells (Duggal and Emerman 2012; Goto et al. 1994). It was later identified as a restriction factor blocking the release of virus particles by linking (tethering) the budding virions to the surface of the virus-producing cell (Neil, Zang, and Bieniasz 2008; Van Damme et al. 2008). The tethering is enabled by the unique topology of tetherin, where two membrane associated domains (a transmembrane N-terminal domain and a glycosylphosphatidylinositol [GPI] C-terminal domain) can span the virus and host membranes. In this way, the tethering effect can extend to multiple enveloped viruses (Mahauad-Fernandez and Okeoma 2016). In response, many viruses developed tetherin antagonists, which at least partially block its antiviral actions (Sauter 2014).

Although tetherin was initially considered to be present only in mammals, it was later identified in other vertebrate genomes, including birds (Blanco-Melo, Venkatesh, and Bieniasz 2016; Krchlíková et al. 2020; Heusinger et al. 2015). Chicken possesses antiviral tetherin, inducible by interferon (IFN). In other birds, intriguing patterns of functional tetherin losses have been described. These include premature stop codon in turkey and pheasant species (Krchlíková et al. 2023), and loss of IFN induction in other galliform birds mediated by mutation of IFN-sensitive response element (ISRE) in tetherin promoter region. Other types of tetherin gene losses are predicted in the genomes of additional avian species (data not shown). Overall, in birds, tetherin seems to have very complex interaction with viruses and any potential, thus far unknown, virus-encoded antagonists. This is in agreement with a strong pattern of diversifying selection detected in intact tetherin orthologs from galliform and passeriform birds (Krchlíková et al. 2020).

To analyze the role of tetherin in *in vivo* infections, we generated tetherin-deficient chicken by using the CRISPR/Cas9 technology employed previously to generate genetically modified chicken (Koslová et al. 2021; Trefil et al. 2017; Koslová et al. 2020). In this work, we describe the generation of tetherin-deficient chicken and initial analysis of their phenotype and susceptibility to virus infection.

## 2. RESULTS

### 2.1 Preparation and Breeding of Tetherin/BST2 KO Chickens

In freshly derived primordial germ cells (PGCs), we introduced indel mutations into the first coding exon of the chicken tetherin gene (Fig. 1A). After nucleofection of the CRISPR/Cas9 construct pX458-tetherin and single cell FACS sorting, we expanded four proliferative clones with indel mutations 5’ to the cleavage site. For further work, we chose one male clone PGC RIR No. 1G6 homozygous for a frame-shifting deletion of 4 nt 5’ to the cleavage site. In order to obtain chickens with a knock out of the tetherin gene, we used orthotopic PGC transplantation into the testes of roosters with suppressed endogenous spermatogenesis (Fig. 1B). Two azoospermic roosters were transplanted and both resumed spermatogenesis after 13 weeks. The presence of 4 nt deletion was detected by PCR in the DNA isolated from the sperm. After insemination, both intramagnal and intravaginal, of four SH hens, we obtained four F_1_ offspring, two males and two females, with the expected tetherin KO +/-genotype. After sexual maturation, F_1_ chickens were mated to produce F_2_ generation with observed tetherin KO -/-, +/-, and +/+ genotypes in a ratio 8 : 56 : 36, respectively. In F_3_ offspring, we observed segregation of tetherin KO genotypes in a ratio of 17 (-/-) : 44(+/-) : 39 (+/+) in correspondence to the expected Mendelian ratio.

**Fig. 1.**
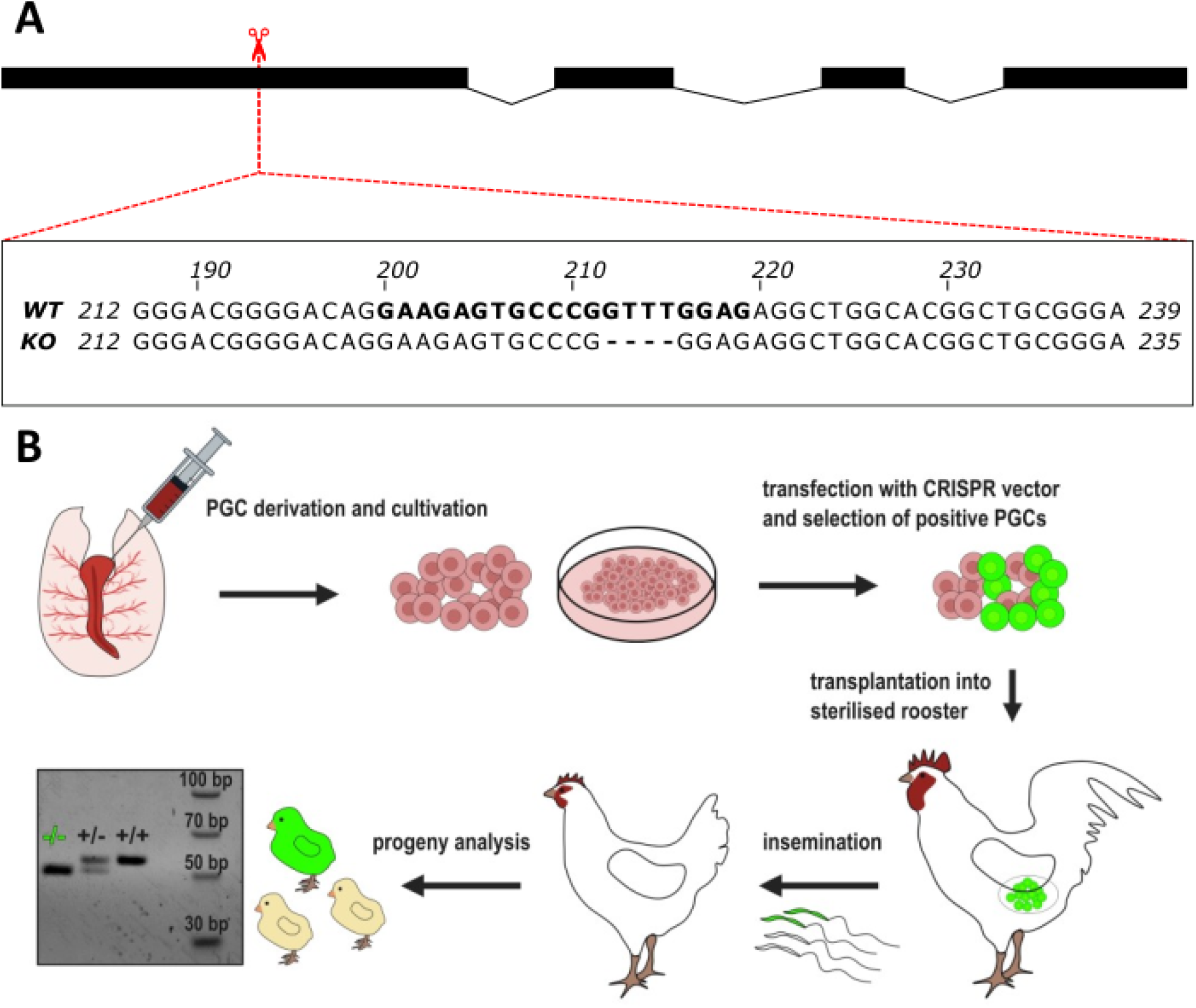
Generation and genotyping of tetherin/BST2 knockout in chicken. (A) Schematic representation of four coding exons of chicken tetherin gene. Position and alignment of sequences around the region targeted by the CRISPR/Cas9 construct are shown (sequences used in the targeting oligonucleotide are highlighted in bold). The 4-bp deletion in the knockout (KO) clone is shown as dashes. (B) Summary of the workflow during generation of the tetherin knockout chicken.

### 2.2 Phenotype and White Blood Cell Count of Tetherin/BST2 Knockout Animals

The tetherin knockout animals do not show any apparent changes in viability or growth compared to wild type siblings (data not shown). The ratio of various genotypes in F_2_ and F_3_ generations approximately complies with expected Mendelian frequencies. In mammals, in addition to antiviral effects, tetherin influences blood cell development and function (Tiwari et al. 2019; Urata et al. 2023; Epeldegui, Blom, and Uittenbogaart 2015; Yoo et al. 2016; Cao et al. 2009). We therefore analyzed the total white blood cell count of knockout animals to characterize their phenotype in more detail. All cell populations were comparable among the three tetherin genotypes (Fig. 2). The only exception was detected in heterophils, whose numbers were significantly higher in tetherin^-/-^ animals.

**Fig. 2.**
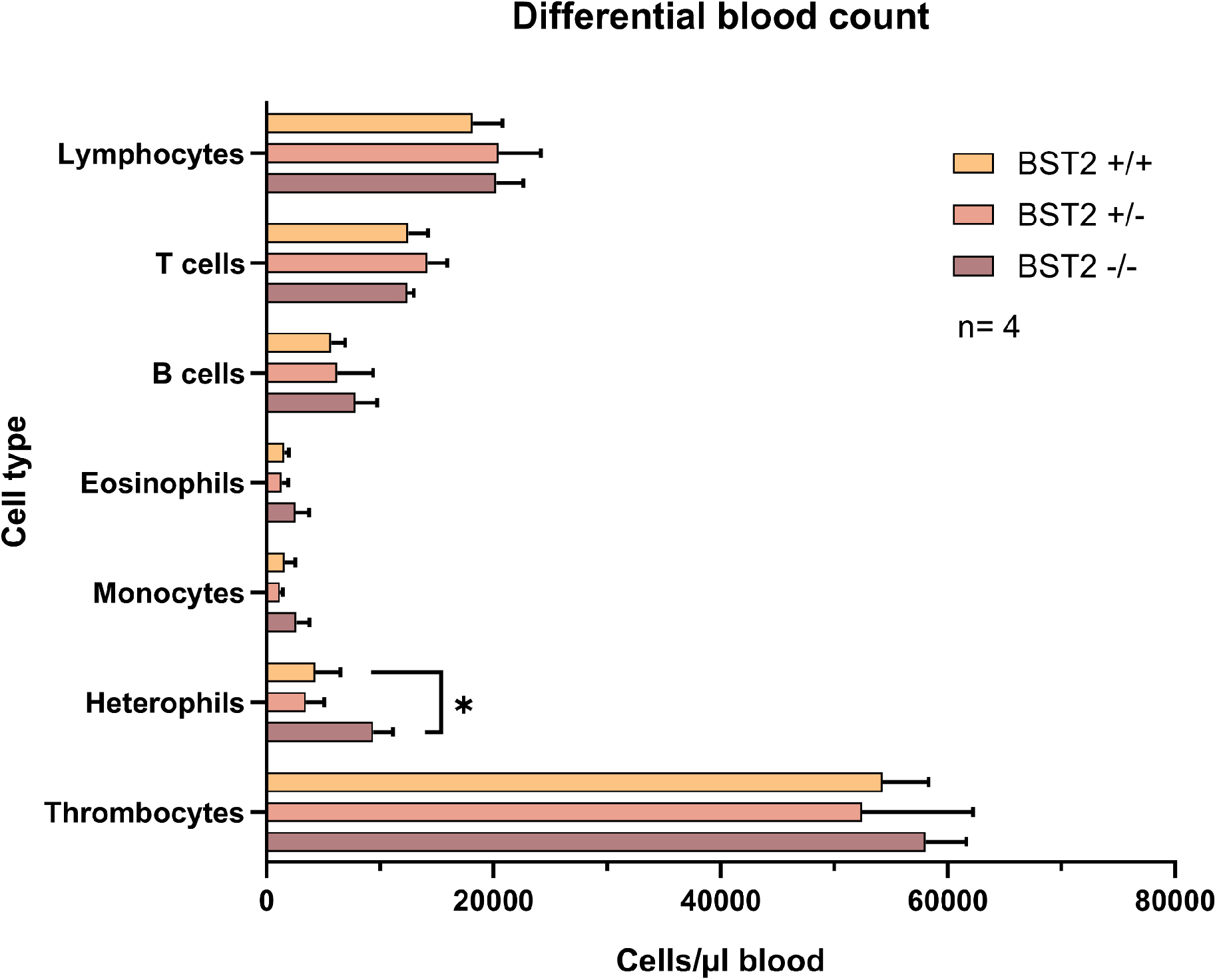
Flow cytometric white blood cell count differential in tetherin knockout chicken. Cell counts for all three genotypes are shown. *; P ≤ 0.05

### 2.3 Susceptibility of tetherin/BST2 Knockout Animals to ASLV Infection

For initial testing of virus susceptibility phenotype, we decided to use avian sarcoma and leukosis virus (ASLV), because this work is well established in our animal facility. We used ASLV-based replication competent vector RCASBP(C)GFP with the envelope (*ENV*) gene of subgroup C virus specificity (Elleder et al. 2005), which encodes a GFP reporter. The outcome of infection was monitored by quantifying viremia in sera of infected animals by quantitative reverse transcriptase PCR (qRT-PCR). Each of the three tetherin genotypes (+/+, +/-, -/-) was represented by 3-7 animals with the age spanning 4-5 weeks. The virus challenge was done by i.v. injection of 10^5^ infectious units RCASBP(C)GFP, the serum was collected from the birds repeatedly at days 6 and 13 post infection (dpi). At both timepoints, the detected viremia was significantly higher in tetherin -/-animals (Fig.3). This is consistent with slightly higher sensitivity of tetherin knockout chicken to ASLV infection.

**Fig. 3.**
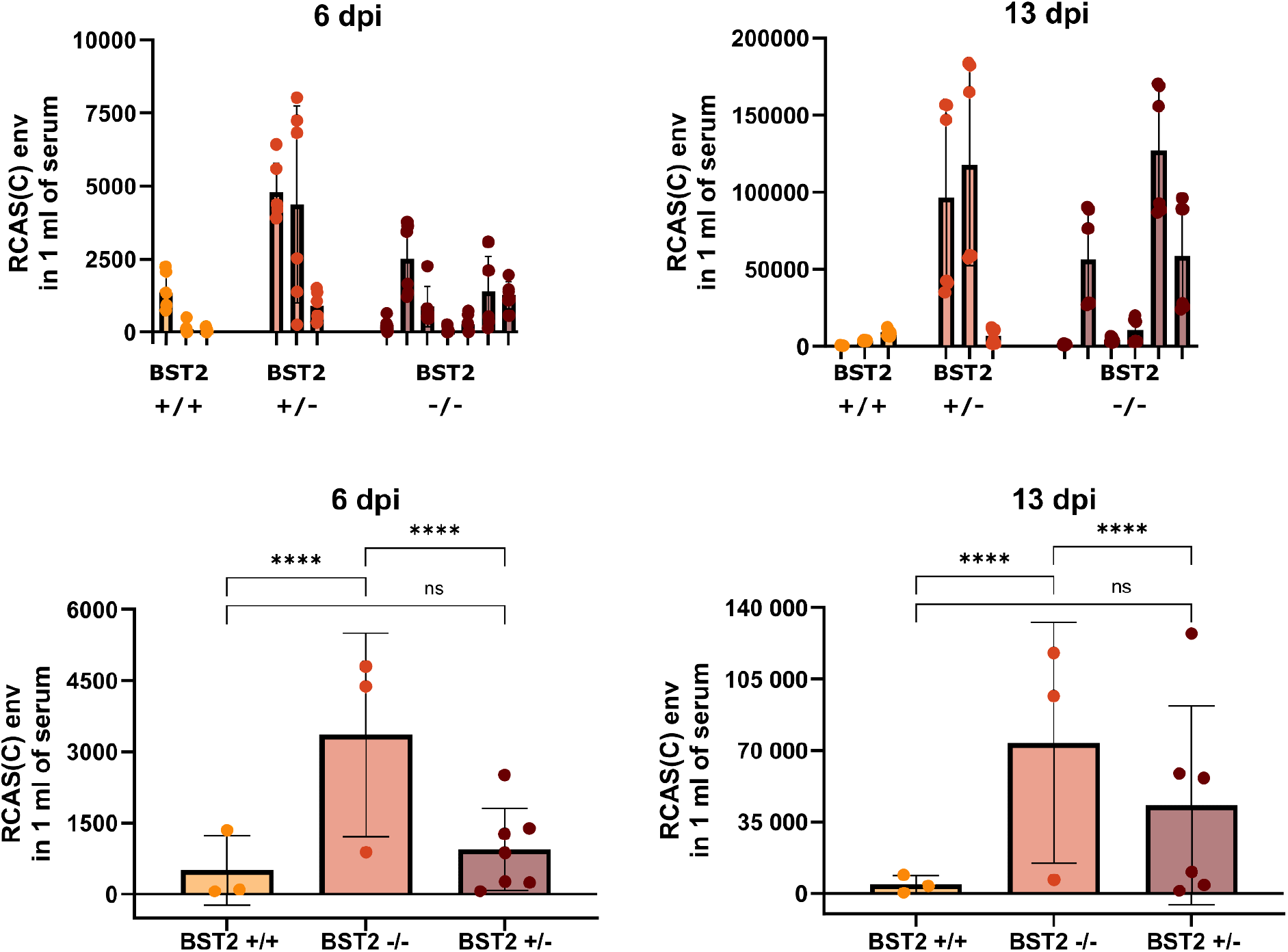
Susceptibility of tetherin/BST2 knockout chicken to in vivo infection with RCASBP(C)GFP vector. Viremia was analyzed by quantification of the *ENV* subgroup C gene in the serum by qPCR. Upper part shows the titer of viral RNA molecules in individual animals, with averages and standard deviation of three technical replicates. Lower part depicts averages and standard deviations for the three genotype groups. ****; P ≤ 0.0001

## 3. DISCUSSION

In this study, we generated a homozygous *in vivo* knockout of the chicken tetherin/BST2 gene. We observed no decrease in viability in the tetherin-deficient animals. This is consistent with the normal viability of tetherin KO in mice (Liberatore and Bieniasz 2011). It is also consistent with the natural absence of functional tetherin in multiple galliform birds (Krchlíková et al. 2023). Multiple physiological functions of tetherin have been described, which are to varying degrees affected by its decrease or absence. These include effects on lipid rafts and platelet receptor signaling, tethering of exosomes and cytokinetic midbody remnants, and organization of the actin cytoskeleton in polarized epithelial cells (Billcliff et al. 2013; Zhao et al. 2021; Edgar et al. 2016; Rollason et al. 2009). Our knockout model can help to dissect these more subtle effects of tetherin in the context of avian cells.

The only significant finding in our phenotypical characterization of tetherin knockout chicken was the increased number of heterophils. These cells are considered the avian equivalent of mammalian neutrophils, and are therefore primary components of innate immunity (Genovese et al. 2013). Heterophils are reactive to microbial stimuli and exhibit chemotaxis, phagocytosis, and production of multiple cytokines. In mammals, tetherin can regulate cytokine production through interaction with ligand ILT7 expressed on plasmacytoid dendritic cells (pDC) (Cao et al. 2009). However, ILT7 is present only in mammals, and the existence of a pDC cell population was only recently suggested in chickens (Wu et al. 2023). Therefore, it is currently not understood how tetherin deficiency could cause the increased heterophil numbers and what are the possible consequences for virus susceptibility.

Tetherin deficiency in mammals does not have major effects on retrovirus replication. The reason is, at least partially, determined by the ability of retroviruses to evade innate immune reaction and IFN induction (Gómez-Lucía et al. 2009; Cingöz and Goff 2019). Without IFN induction, tetherin expression is restricted to several cell types (Goto et al. 1994). Consequently, the replication and disease progression of Moloney murine leukemia virus (Mo-MLV) was similar in wild-type and tetherin-deficitent mice (Liberatore and Bieniasz 2011). External IFN induction was needed to allow tetherin induction and to reveal its antiviral activity. Later work in the mammalian system showed tetherin influence on cell-mediated antiviral responses and sometimes paradoxical positive effects on virus replication (Li et al. 2014; Swiecki et al. 2012). In this study, we detected higher ASLV viremia in tetherin knockout chickens, without externally added IFN. One possibility is that constitutive expression of tetherin is higher in chickens than in mammals and such endogenous levels are sufficient for antiviral activity. Indeed, analysis of publicly available RNAseq data from various chicken tissues shows substantial tetherin mRNA expression in several organs (Fig. 4). Further studies, including generation of antibodies against chicken tetherin, are needed to characterize its *in vivo* expression profile and response to retrovirus infection (work in progress). It will also be interesting to assess the susceptibility of tetherin KO chicken to additional enveloped avian viruses, which are potential targets of tetherin. Some of these viruses, including Newcastle disease virus (NDV), avian influenza viruses (AIV) and Usutu virus, are more potent IFN inducers than retroviruses, and might therefore be even more affected by tetherin absence.

**Fig. 4.**
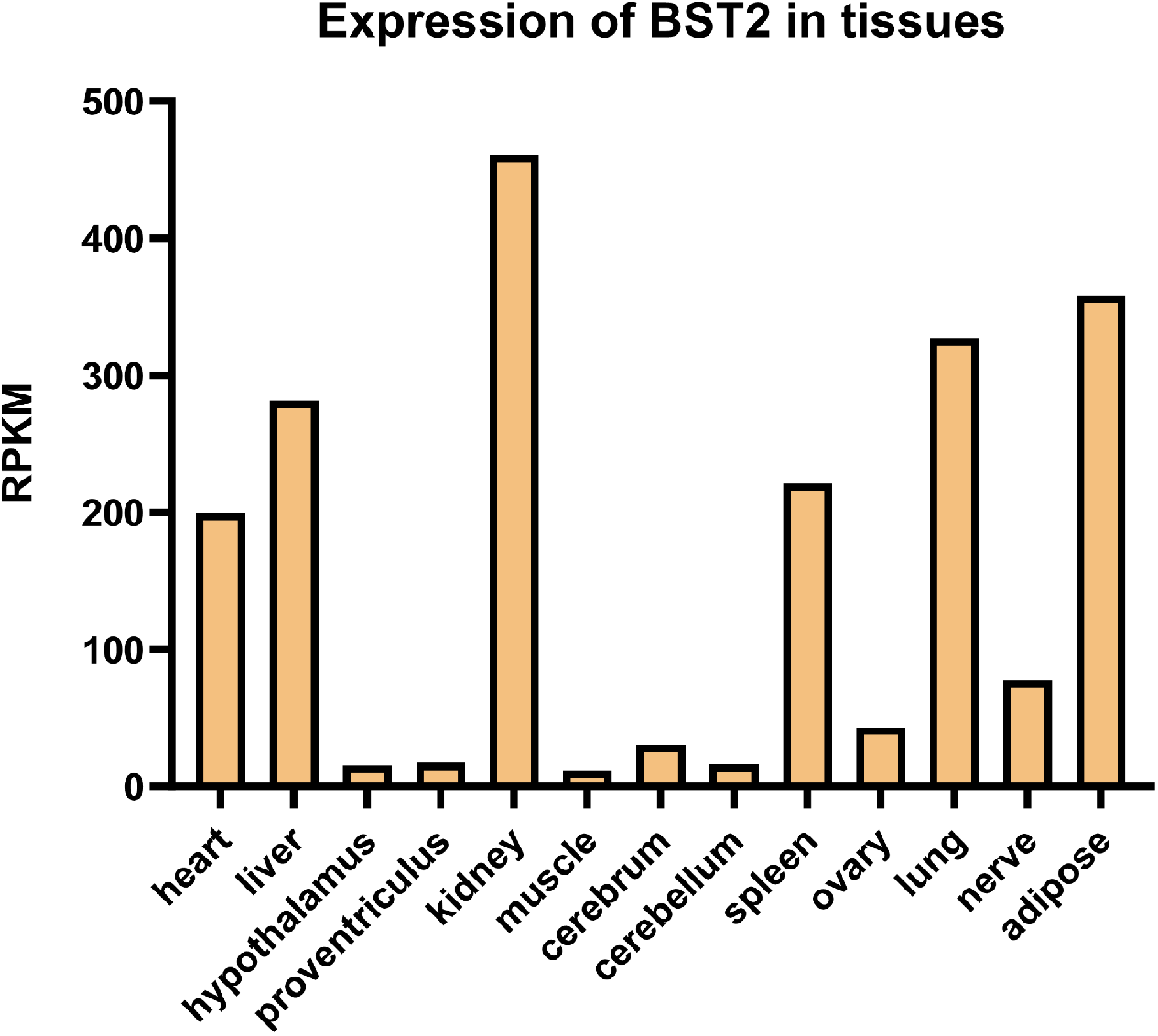
Expression pattern of tetherin/BST2 in various chicken tissues. The expression values were obtained from NCBI Sequence Read Archive (SRA) data PRJEB4677 as described in Methods and expressed in relative expression units (RPKM, Reads per kilobase per million mapped reads).

In summary, we introduce a new *in vivo* knockout model of the chicken antiviral gene tetherin/BST2. Transgenesis in birds remains challenging, but there were major advances in the application of gene editing technology in chicken, which represents an important model in the fields of immunology and infectious disease (Woodcock, Idoko-Akoh, and McGrew 2017; Sid and Schusser 2018; Khwatenge and Nahashon 2021). As in other animal models, the CRISPR/Cas9 system is revolutionizing the possibilities of gene modification. The availability of tetherin knockout animals should facilitate the *in vivo* studies of the complex antiviral defense in birds.

## 4. METHODS

### Ethical Statement

In this work, we conducted all experiments and procedures in accordance with the Czech legislation for animal handling and welfare. The Animal Commodities Department of the Ministry of Agriculture of the Czech Republic approved all animal experiments described in this study (approval no. 65823/2019-MZE-18134).

### Experimental Chickens

Embryos of the commercial chicken breed RIR D159 (Líheň Studenec, Czech Republic) were used as a source of PGCs and hybrid roosters (♂ CC21 × ♀ L15) were used as recipients in orthotopic PGC transplantation. Hens of the breed SH were used for insemination. Inbred lines CC21, L15, and outbred population SH were maintained at the Institute of Molecular Genetics, Czech Academy of Sciences, Prague, Czech Republic (Plachý 2000). Standard husbandry conditions applied in this work included 16 h light/8 h dark cycle, food/water provided ad libitum, and housing the birds in either deep litter individual cages (recipient roosters) or battery cages (inseminated hens). The laid eggs were incubated in a forced air incubator (BIOS MIDI).

### PGC Cultivation and Editing of Tetherin/BST2 Gene

We derived PGCs from the blood aspirations of RIR D159 embryos at 2.5 days of incubation, which corresponds to the Hamburger and Hamilton stage 16. Specifically, we transferred 5 μL of blood from the dorsal aorta on a 48-well plate with 150 μL of Avian KO-DMEM (Thermo Fisher Scientific, Waltham, MA, USA) supplemented with growth factors, as described previously (Mucksová et al. 2019). We cultured the PGCs for 3 weeks, expanded them up to 10^5^ cells, and determined their sex by detection of the W chromosome. Oligonucleotide primers and cycling conditions used for W detection were as described in Mucksová et al. (Mucksová et al. 2019). DNA for W amplification was isolated from PGC aliquots using the PureGene kit (Qiagen, Hilden, Germany). Two proliferative male PGC lines (RIR PGC 4 and RIR PGC 7) were selected for the tetherin knock-out at the age of 42 d in culture.

We used the CRISPR/Cas9 construct pX458-tetherin already adjusted for targeting the first exon of the chicken tetherin gene (Krchlíková et al. 2020). Suspension of 2.5 × 10^5^ PGCs in Nucleofector Solution V (Lonza) was mixed with 10 μg of pX458-tetherin in total volume of 100 μL and nucleofection was performed with the AMAXA nucleofector (Lonza) using the A-27 program. GFP-positive cells with the highest fluorescence intensity (top 5%) were single-cell-sorted 3 d post nucleofection and expanded for two weeks. The resulting clones were characterized as to the hit of the target sequence of the chicken tetherin gene; oligonucleotide primers and PCR conditions used for tetherin characterization are described in Krchlíková et al. (Krchlíková et al. 2020).

### Sterilization of Recipient Roosters and Orthotopic PGC Transplantation

We used orthotopic PGC transplantation into roosters with suppressed endogenous spermatogenesis in order to obtain founder animals transducing the edited tetherin allele in offspring F1 generation. The procedures of rooster irradiation and PGC transplantation were described previously (Koslová et al. 2020; Trefil et al. 2017; Mucksová et al. 2019). Briefly, hybrids of inbred lines CC21I × L15 at the age of 8 months were irradiated with five doses of 8 Gy over two weeks using the Terabalt radiation unit (UJP Prague) with Co^60^ as a source of gamma rays. Subsequent decline of endogenous spermatogenesis was monitored in semen samples collected by dorsoabdominal massage. Azoospermic recipient roosters were anesthetized and bilaterally injected through tunica albuginea with a dose of 7.4 × 10^6^ PGCs in 250 μL of culture medium. Anesthesia was performed by intramuscular injection of 15 mg/kg ketamine (Narkamon, Bioveta, Czech Republic) and 4 mg/kg xylazine (Rometar, Bioveta, Czech Republic). We did not encounter any mortality or side effects associated with irradiation, anesthesia, or transplantation surgery.

The appearance of PGC-derived spermatogenesis was monitored in semen samples collected by dorsoabdominal massage starting from the third week after transplantation. The threshold for intramagnal insemination was the semen concentration of 10^4^ sperms per mL, and 0.1 to 0.4 mL of undiluted semen was used as the insemination dose.

### Genotyping PCR for tetherin locus

Short genomic sequence containing the CRISPR/Cas9-induced deletion site in the tetherin locus was PCR-amplified with primers 5’ TGGGGACGGGGACAGGAAGA and 5’ TCCCGCAGCCGTGCCA from blood lysates. Deletion of 4 nucleotides was then directly checked on 4% low melting MetaPhor® agarose gel (Lonza) and in some cases verified by sequencing.

### Flow cytometry method for total white blood cell count

The differential white blood cell count was done as described in Seliger et al. (Seliger et al. 2012). Briefly, EDTA blood samples of 3-6 weeks old animals were stabilized by the addition of TransFix Reagent (R) (Cytomark, Buckingham, UK); 24 h later,1μl of blood was diluted 1:50 and stained with an antibody mixture of anti-chicken CD45-APC (clone 16-6; 1:2.000), anti-chicken CD4 FITC (clone CT4; 1:2.500), anti-chicken CD8-FITC (clone CT8 1:2.500), anti-chicken TCRγδ-FITC (clone TCR1; 1:300), Bu1-PerCP-Cy5.5 (clone AV20, 1:800), K1-RPE (clone K1, 1:1.000) and anti-chicken MRC1L-B-RPE (clone KulO1, 1:2.000) using the described no-lyse no-wash single-step one-tube technique. CD4-FITC, CD8-FITC, TCRγδ-FITC and MRC1L-B-RPE were obtained from Southern Biotech (Birmingham, USA). CD45-APC and Bu1-PerCP-Cy5.5 were generated by conjugation of the antibody with the respective Lynx Rapid conjugation kit (Bio-Rad, Feldkirchen, Germany). Samples were analyzed on a FACS Canto II and FlowJo 10.9 software (BD, Heidelberg, Germany).

### In vivo infection of chicken with ASLV vector and quantitative PCR

Animals of all three tetherin genotype groups were infected by i.v. injection of RCAS-C-GFP, with dose 10^5^ infectious units (IU). 6 and 13 dpi blood samples were obtained and coagulated into blood sera. For viral RNA isolation from the chicken sera, QIAamp Viral RNA Mini Kit (Qiagen) was used. Two independent isolations were performed according to the manufacturer’s protocol from 140 μL and 280 μL of sera respectively. Then we used ProtoScript® II First Strand cDNA Synthesis Kit (NEB) and 3′-RACE CDS primer (ClonTech) for cDNA synthesis from 5,4 μL uf viral RNA. Next, qPCR was performed using PowerTrack SYBR Green Master Mix (Thermofisher) with primers for ASLV-C env (5’-TGTGTATATTTCGCCCCAAGG, 5’-CCTCTGCACCAGGTTTCAGG). We used serial dilution of RCASBP-C plasmids to generate the standard curve for absolute gene quantification. Reaction was run in the QuantStudio™ 5 Real-Time PCR System (Thermofisher) with the following protocol: one cycle of 8 min at 95°C then 40 cycles of 15 s at 95°C, 25 s at 60°C, and 35 s at 72°C and final polymerization step at 72°C for 10 min.

### Analysis of publicly available RNAseq data

To obtain the *in silico* mRNA expression pattern of tetherin, its coding sequences were used in blast searches against the SRA data PRJEB4677. Number of blast hits from RNAseq datasets of individual chicken tissues were recalculated to obtain relative measures of mRNA expression levels (RPKM; Reads per kilobase per million mapped reads).

### Statistical tests

For statistical evaluation of the differences between WT and KO animals, unpaired Student’s t-test was used.

## 5. ACKNOWLEDGEMENTS

This work was funded by grant 20-22063S from the Czech Science Foundation. D.E. and J.H. were further supported by the project National Institute of virology and bacteriology (Programme EXCELES, No. LX22NPO5103) funded by the European Union—Next Generation EU.

